# Target Capture Sequencing of SARS-CoV-2 Genomes Using the ONETest Coronaviruses Plus

**DOI:** 10.1101/2021.03.25.437083

**Authors:** Shing H. Zhan, Sepideh M. Alamouti, Brian S. Kwok, Meng-Hsun Lee, Jaswinder Khattra, Habib Daneshpajouh, Herbert J. Houck, Kenneth H. Rand

## Abstract

**Background:** Genomic sequencing is important to track and monitor genetic changes in SARS-CoV-2. We introduce a target capture next-generation sequencing methodology, the ONETest Coronaviruses Plus, to sequence SARS-CoV-2 genomes and select genes of other respiratory viruses simultaneously.

**Methods:** We applied the ONETest on 70 respiratory samples (collected in Florida, USA between May and July, 2020), in which SARS-CoV-2 had been detected by a qualitative PCR assay. For 48 (69%) of the samples, we also applied the ARTIC protocol for Illumina sequencing. All the libraries were sequenced as 2×150 nucleotide reads on an Illumina instrument. The ONETest data were analyzed using an in-house pipeline and the ARTIC data using a published pipeline to produce consensus SARS-CoV-2 genome sequences, to which lineages were assigned using *pangolin*.

**Results:** Of the 70 ONETest libraries, 45 (64%) had a complete or near-complete SARS-CoV-2 genome sequence (> 29,000 bases and with > 90% of its bases covered by at least 10 reads). Of the 48 ARTIC libraries, 25 (52%) had a complete or near-complete SARS-CoV-2 genome sequence.

In 24 out of 34 (71%) samples in which both the ONETest and ARTIC sequences were complete or near-complete and in which lineage could be assigned to both the ONETest and ARTIC sequences, the SARS-CoV-2 lineage identified was the same.

**Conclusions:** The ONETest can be used to sequence the SARS-CoV-2 genomes in archived samples and thereby enable detection of circulating and emerging SARS-CoV-2 variants. Target capture approaches, such as the ONETest, are less prone to loss of sequence coverage probably due to amplicon dropouts encountered in amplicon approaches, such as ARTIC. With its added value of characterizing other major respiratory pathogens, although not assessed in this study, the ONETest can help to better understand the epidemiology of infectious respiratory disease in the post COVID-19 era.

## INTRODUCTION

SARS-CoV-2 genome sequencing is widely achieved using the amplicon next-generation sequencing (NGS) ARTIC methodology ^1^. Because of its ease of use and low cost of sequencing, ARTIC has become the method of choice among many laboratories. Notwithstanding its advantages, the ARTIC PCR primer set needs to be maintained and updated due to amplicon dropouts ^1^, which may be caused by primer interactions ^2^ and mutations at primer binding sites ^3^. Without continual upkeep, amplicon sequencing may yield incomplete SARS-CoV-2 genome sequences and therefore create a loss of valuable genetic information. This could weaken our vigilance towards SARS-CoV-2 mutations, which may impact our diagnostic, therapeutic, and vaccination efforts ^4^, and SARS-CoV-2 lineages, especially variants of concern such as B.1.1.7 and B.1.135 that may enhance the virus’ transmissibility or lethality ^5,6^

Alternatively, SARS-CoV-2 genome sequencing can be accomplished using probe-based liquid-phase hybridization followed by NGS ^3,7,8^. A major appeal of target capture NGS methodologies is its capacity to enrich samples for a practically limitless repertoire of genetic loci without needing to constantly update the primers and or deal with multiplexing issues encountered with amplicon-based approaches. Indeed, virome target capture NGS methodologies have been developed (e.g., ^9,10^). Another advantage is that target capture NGS approaches perform better than amplicon NGS approaches in degraded samples (e.g., archived FFPE samples ^11^). A validated target capture NGS solution with end-to-end automation for concurrent detection and sequence characterization of SARS-CoV-2 and other common respiratory pathogens can be a powerful tool for genomic surveillance of respiratory infectious disease in the post COVID-19 era and can play a crucial role in timely generation and dissemination of genomic data.

The ONETest^™^ is a pre-commercial target capture NGS platform developed by Fusion Genomics Corp. (Burnaby, BC, Canada). The platform offers a sequencer-agnostic end-to-end NGS workflow that includes library preparation, probe-based liquid phase hybridization, and cloud-based bioinformatics analysis. The ONETest^™^ Coronaviruses Plus (http://www.fusiongenomics.com/onetestplatform/coronavirusesplus/), based on the ONETest^™^ platform, has been demonstrated to enrich samples for select genetic loci of various respiratory viruses (e.g., influenza A viruses) in a separate study (*in preparation*). Furthermore, the ONETest^™^ EnviroScreen, also based on the ONETest^™^ platform, has been shown to detect diverse subtypes of avian influenza viruses in wetland sediments ^12^.

To capture the full-length genome of SARS-CoV-2, we have expanded the probe design of the ONETest Coronaviruses Plus. Here, using the updated ONETest, we sequenced the SARS-CoV-2 genomes in 70 retrospectively selected samples, which were initially tested at the University of Florida (UF) Health Shands Hospital Clinical Laboratory during the COVID-19 pandemic in 2020. We also processed a subset of them (n = 48) using the ARTIC protocol for Illumina sequencing. These data allowed us to demonstrate the ability of the ONETest to determine the genome sequences of SARS-CoV-2 from respiratory samples.

## RESULTS

### ONETest yields complete or near-complete SARS-CoV-2 genome more often than ARTIC

The ONETest libraries of the 70 samples had a total of ~186 million paired-end reads, and each of the libraries had ~2.66 million paired-end reads on average (range, ~0.45 to ~6.14 million) (**Table S1**). This per-sample amount of sequencing is comparable to that used in a study ^3^ evaluating another target capture product (7.4 million 1×100 nt filtered reads per sample). Of the 70 ONETest libraries, 45 (64%) had a complete or near-complete SARS-CoV-2 genome sequence that was > 29,000 nucleotides (nt) long and had > 90% well covered bases (specifically, ≥ 10x depth). Even after sub-sampling, the ONETest libraries had a complete or near-complete genome sequence for 43 (61%) of the samples. Additionally, we processed 48 (69%) of the 70 samples using ARTIC. The ARTIC libraries had a total of ~30 million paired-end reads, and each of the libraries had ~0.63 million paired-end reads on average (range, ~0.20 to ~2.1 million) (**Table S1**). This amount of sequencing is comparable to that in the ARTIC experiments performed by other groups (**Figure S1**). Of the 48 ARTIC libraries, 25 (52%) had a complete or near-complete SARS-CoV-2 genome sequence.

When considering the 48 samples for which both ONETest and ARTIC libraries were made, the mean percent poorly covered bases in the ONETest sequences was 23% (range, 0% to 100%), whereas that in the ARTIC sequences was 25% (range, 3% to 99%) (**Table S1**). For 34 (71%) of the samples, there was sufficient sequence information in both the ONETest and ARTIC libraries so that lineage could be assigned to both the ONETest and ARTIC sequences using *pangolin* (see below). We focused on these lineage-assigned matched ONETest and ARTIC library pairs to compare the genome sequences from the two methodologies.

In the matched ONETest and ARTIC library pairs, there were fewer poorly covered bases (< 10x depth) across the SARS-CoV-2 genome in the ONETest libraries than in the ARTIC libraries (**Figure 1**; **Figure S2**). Some of this difference may be explained by the fact that the ONETest libraries were sequenced deeper than the ARTIC libraries (almost four times deeper on average). However, a sub-sampling analysis indicated that even at similar sequencing depths, the ONETest libraries yielded better sequence coverage than the ARTIC libraries (**Figure S3**).

**Figure 1.**
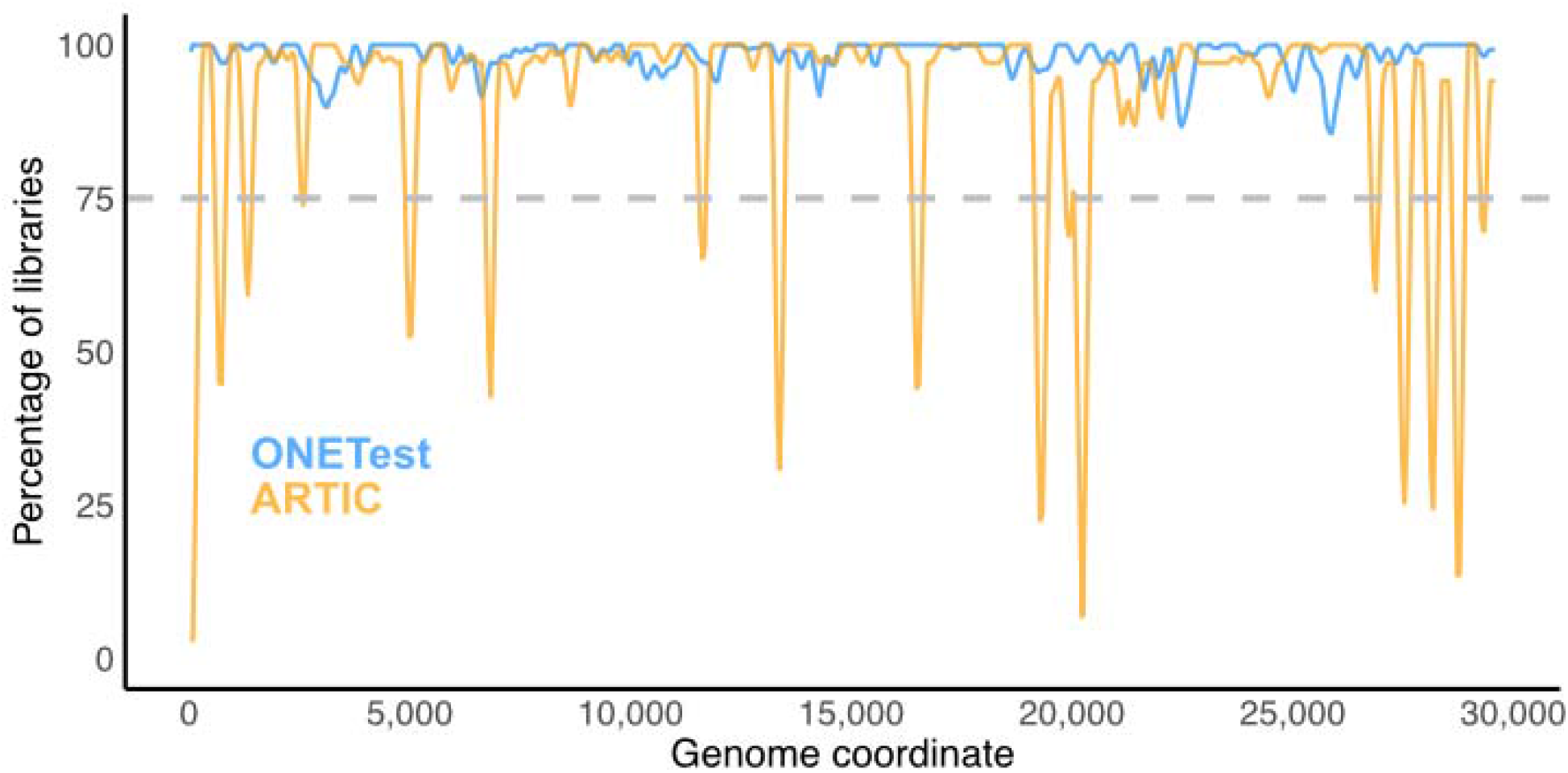
Aggregate summary of sequence coverage over the SARS-CoV-2 genome in the ONETest and ARTIC libraries from the samples examined in this study. Here, we considered only the 34 samples for which lineage could be assigned to both its ONETest and ARTIC sequences using *pangolin*. For each position in the SARS-CoV-2 reference sequence targeted by the ARTIC PCR primers (MN996528.1: 30 to 29,866), we computed the percentage of samples in which its depth of coverage was ≥ 10 (excluding duplicates for the ONETest libraries and including duplicates for the ARTIC libraries). This percentage was averaged across the positions of each 200 nt partially overlapping window across the genome (skip size of 50 nt). Poorly covered regions in the SARS-CoV-2 genome appear as troughs below the dashed line.

### Regions with poorer sequence coverage in the ARTIC libraries than the ONETest libraries

While there were several regions of the SARS-CoV-2 genome in the ARTIC libraries that had poor sequence coverage compared to the ONETest libraries, we closely examined one region that had particularly poor sequence coverage in the ARTIC libraries (**Figure 1**). We observed that depth of coverage was generally poor in the ~19,900-20,500 region of the SARS-CoV-2 genome in the ARTIC libraries (**Figure 1**). This region is targeted by the ARTIC primer pairs 66_LEFT/66_RIGHT (pool 2, MN908947.3: 19,844-20,255) and 67_LEFT/67_RIGHT (pool 1, MN908947.3: 20,172-20,572). In contrast, the ~19,900-20,500 region was well covered overall in the ONETest libraries (**Figure 1**). For example, depth of coverage across the SARS-CoV-2 genome in the ARTIC library of sample 27 was high (mean, 3,937x), except in that region amplified by the two primer pairs (visualized using IGV ^13^ in **Figure S4**); on the other hand, the ONETest library of sample 27 had high depth of coverage across the virus’ genome (mean, 10,354x with duplicate reads and 1,237x without duplicate reads), even in the region targeted by those two problematic ARTIC PCR primer pairs (**Figure S4**).

### Difference in sequence coverage between samples positive for three genes by PCR and samples positive for one or two genes by PCR

In some ONETest and ARTIC libraries, incomplete SARS-CoV-2 genome sequences might have arisen from low-titer samples. Because we used a qualitative PCR assay, we did not have quantitative estimates of viral titer in the samples. Instead, we considered the samples in which three SARS-CoV-2 genes (N, RdRp, and E) were detected by the PCR assay to be of relatively high titer (although some might be of low titer), whereas the samples in which one or two genes (N only, or both N and RdRp) were detected to be of relatively low titer (although some might be of high titer). We noticed that the ONETest and ARTIC libraries from the low-titer samples yielded less complete SARS-CoV-2 genome sequences than the libraries from the high-titer samples. The ONETest sequences from the low-titer samples had more poorly covered bases (mean ± standard deviation; 75% ± 28%) than those from the high-titer samples (2% ± 6%) (p < 0.001, Wilcoxon’s test; including only the ONETest libraries with the matched ARTIC libraries). In line with this observation, the ARTIC sequences from the low-titer samples had more poorly covered bases (60% ± 34%) than those from the high-titer samples (12% ± 13%) (p < 0.001; Wilcoxon’s test).

### ONETest and ARTIC determined SARS-CoV-2 genome sequences with concordant lineage assignments

For 34 samples, the consensus sequences from both the ONETest and ARTIC libraries could be assigned to a SARS-CoV-2 lineage using *pangolin*. In 24 (71%) of these samples, the lineage assignment was identical for the ONETest and ARTIC libraries (e.g., in sample 50, both the ONETest and ARTIC sequences were assigned to B.1.509). In the other 10 samples, the lineage assignment was nevertheless in the same major lineage (e.g., in sample 46, both the ONETest and ARTIC sequences were assigned to the B.1 lineage rather than the A.1 lineage). These differences in lineage assignment likely stemmed from differences in sequence coverage between the ONETest and ARTIC libraries. In the 10 samples, the mean difference in percent poorly covered bases between the ARTIC and ONETest sequences was 5.3%.

### SARS-CoV-2 lineages detected in the ONETest libraries

Of the 70 samples sequenced in this study using the ONETest, 45 had a complete or near-complete SARS-CoV-2 genome sequence. We found 15 distinct SARS-CoV-2 lineage assignments to the ONETest sequences of the samples (**Figure 2**).

**Figure 2.**
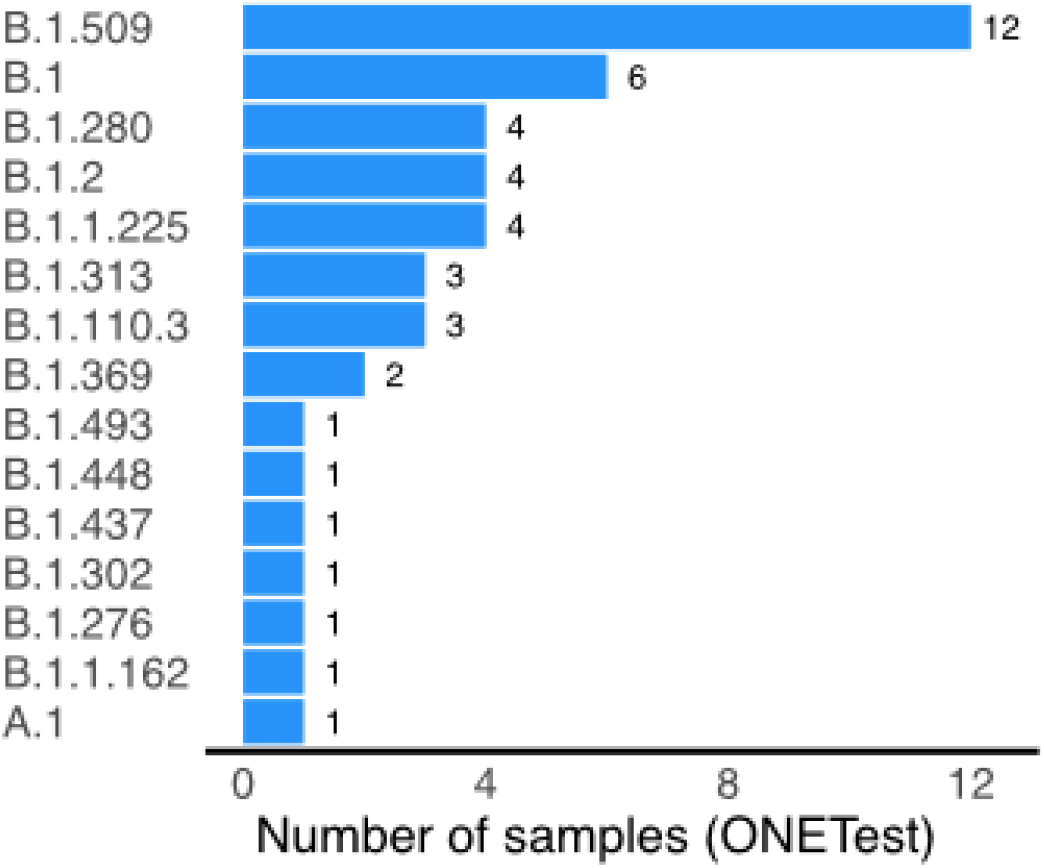
SARS-CoV-2 lineages identified in the samples examined in this study using the ONETest. Lineage was assigned to the complete or near-complete SARS-CoV-2 genome sequences from the ONETest libraries of 45 samples.

## DISCUSSION

Vaccines against SARS-CoV-2 are presently being administered around the globe, but we have yet to see how effectively the vaccines will protect our populations from the new variants of concerns. Having multiple technologies in our SARS-CoV-2 genome sequencing toolbox should help to heighten our vigilance towards new SARS-CoV-2 variants that may escape our vaccines. Here, we propose the ONETest target capture NGS methodology to sequence SARS-CoV-2 genomes to aid in efforts to track SARS-CoV-2 variants.

Using the ONETest and ARTIC, we sequenced SARS-CoV-2 genomes from archived samples in which SARS-CoV-2 had been detected by a FDA EUA qualitative PCR assay. Our data demonstrate that the ONETest can yield complete SARS-CoV-2 genome sequences more often than ARTIC (64% versus 52%). While relatively shallow sequencing of the ARTIC libraries may account for some of the other poorly covered regions, a sub-sampling analysis indicates that the ONETest produces complete genome sequences more often than ARTIC even at about one fourth the amount of sequencing on average. Nonetheless, there are consistently poorly covered regions in the SARS-CoV-2 genome across the ARTIC libraries. In particular, the ~19,900-20,500 SARS-CoV-2 genome region targeted by two ARTIC PCR primer pairs (e.g., sample 27) is poorly covered in many ARTIC libraries, even though other genomic regions in the same libraries are well covered. As shown by an analysis of the SARS-CoV-2 genome sequences deposited in GISAID ^14^, many publicly available sequences contain problematic regions (i.e., contiguous stretches of 200 Ns) around the 20,000th nucleotide position. Many of the genome sequences were produced using an amplicon NGS methodology, in particular ARTIC. Furthermore, by comparing the lineage assignments of the ONETest and ARTIC sequences, which are generally concordant, we show that the ONETest can provide quality genome sequences to study the evolution and epidemiology of SARS-CoV-2.

In this study, we did not have quantitative estimates of viral load (e.g., cycle threshold values from a quantitative PCR assay) for the samples examined here to directly observe the effect of viral load on the quality of the consensus sequences. By using the number of target genes detected by a PCR assay (three genes versus one or two genes) as a proxy instead, we find that the ONETest and ARTIC consensus sequences are of higher quality in the samples positive for three genes, suggesting that the partial genome sequences in about 40% of the ONETest libraries and about 50% of the ARTIC libraries resulted from low viral titer.

Target capture NGS methodologies, such as the ONETest, should be able to detect mutations that can impact the performance of amplicon NGS methodologies, such as ARTIC. Kim et al. ^3^ showed a case in which target capture NGS detected a large 382 nt deletion in the ORF8 gene of SARS-CoV-2 that ablated sequence coverage in four contiguous genes (ORF3a, E, M, and ORF6) in the ARTIC library due to PCR amplification failure. Although we did not encounter such a dramatic case in this study, we anticipate that as we sequence more samples using the ONETest, the ONETest will detect large deletions in the SARS-CoV-2 genome that could severely reduce sequence coverage when using amplicon NGS methodologies. This advantage of target capture NGS approaches is important as new SARS-CoV-2 genetic mutations of unpredictable nature continue to emerge.

Our data show the ability of the ONETest to determine the genome sequences of SARS-CoV-2 in respiratory samples. Importantly, our data indicate that the ONETest is less prone to loss of sequence coverage that may be caused by poor or failed target binding (e.g., the amplicon dropouts in the ARTIC libraries shown here and in studies by other groups), which can ultimately result in inaccurate SARS-CoV-2 genotyping and lineage identification. The added value of the ONETest to characterize multiple respiratory pathogens, although not assessed in this study, would help us to better understand the epidemiology of respiratory pathogens in the post COVID-19 era. Furthermore, Fusion Genomics Corp., at the time of this writing, is validating a fully automated ONETest workflow that allows for flexible sample batching (i.e., as few as eight libraries to as many as 384 libraries per sequencing run).

## MATERIALS AND METHODS

### Ethics review

Approval for this study was obtained from the University of Florida Institutional Review Board (IRB202001328).

### Respiratory samples

Nasopharyngeal (NP) swabs (n = 61) and endotracheal aspirates (n = 9) were collected from patients, who had respiratory illness and were suspected to have COVID-19, at UF Health Shands Hospital in May (n = 31) and in July (n = 39), 2020. Among the patients, 30 (43%) were male and 40 (57%) were female. The mean age of the patients (± standard deviation) was 46.1 (± 19.8) years (range, 5 to 102 years; interquartile range, 27.8 to 54.0 years). Three patients had two separate samples collected seven to 12 days apart; one patient had four samples, two samples collected in May (one NP swab and one endotracheal aspirate on the same day) and two samples collected in July that were duplicate samples. The samples were initially tested for SARS-CoV-2 using a FDA Emergency Use Authorization qualitative PCR assay (GeneFinder^™^ COVID-19 Plus RealAmp Kit from OSANG Healthcare Co. Ltd., South Korea), which targets the RdRp, N, and E genes. We retrospectively selected 70 samples in which SARS-CoV-2 had been detected by the PCR assay.

### RNA extraction

Nucleic acids were isolated from 200 μL of the samples and eluted in 100 μL, of which 10 μL was tested for SARS-CoV-2 by the ELlTe InGenius^®^ platform (ELITechGroup, Puteaux, France) using the GeneFinder^™^ COVID-19 Plus RealAmp Kit, as per the manufacturer’s instructions. The remaining 90 μL of de-identified RNA extracts were then shipped to Fusion Genomics Corp. (Burnaby, BC, Canada). Each RNA extract was treated with DNAse (MilliporeSigma Canada, Ontario) and partitioned into two aliquots. One aliquot was processed using the ARTIC protocol and the other using the ONETest protocol.

### ARTIC protocol

We processed 2 μL of RNA extract from each sample using the ARTIC Illumina protocol (https://www.protocols.io/view/covid-19-artic-v3-illumina-library-construction-an-bibtkann). This protocol utilizes two pools of ARTIC V3 primer pairs to amplify 98 ~400 nt partially overlapping regions that tile the entire SARS-CoV-2 genome (https://github.com/artic-network/artic-ncov2019/blob/master/primer_schemes/nCoV-2019/V3/), which were ordered from Sigma-Aldrich (Oakville, ON, Canada). Libraries were constructed using TruSeq Nano from Illumina Inc. (San Diego, CA, USA), as per the manufacturer’s instructions. Libraries were normalized, pooled together, and sequenced as 2×150 nt reads on an Illumina NextSeq 500 instrument (San Diego, CA, USA). Reads from these libraries were analyzed using a bioinformatics pipeline (v1.3.0; https://github.com/connor-lab/ncov2019-artic-nf) that automates the ARTIC data analysis protocol for Illumina reads (https://artic.network/ncov-2019/ncov2019-bioinformatics-sop.html), which utilizes *bwa mem* ^15^, *samtools* ^16^, and *iVar* ^17^.

### ONETest: probe design

We have expanded the ONETest probe set (QuantumProbes^™^; http://www.fusiongenomics.com/onetestplatform/), which originally targets non-SARS-CoV-2 respiratory pathogens, to capture the entire SARS-CoV-2 genome based on the Wuhan-Hu-1 reference sequence (NC_045512.2). Additionally, we designed probes to capture the nucleotide variants frequently observed in SARS-CoV-2 genomes (> 1%; retrieved from NCBI GenBank in July, 2020) and to cover the GC-poor regions (< 35% GC) of the virus’ genome.

### ONETest: library preparation, target capture, and NGS

Next, we processed 11 μL of RNA extract from each sample using the ONETest protocol. RNA extracts were then treated with deoxyribonuclease from Sigma-Aldrich (Oakville, ON, Canada). Target-enriched Illumina-compatible libraries were prepared from RNA using the ONETest kit from Fusion Genomics Corp. (Burnaby, BC, Canada). Total RNA was subject to rRNA and mRNA removal using biotin-labeled depletion probes captured via magnetic streptavidin-coated beads. Cleaned RNA was then reverse transcribed using random primers with adapters, and the resulting cDNA was fragmented. Whole transcriptome amplification was then performed, and cDNA was ligated with Illumina-compatible indexed adapters, according to the manufacturer’s instructions. The indexed libraries were mixed with Illumina adapter-specific blocking reagents, human Cot-1 placental DNA from Sigma-Aldrich (Oakville, ON, Canada), and target-specific biotin-labeled probes in hybridization solution. Hybridization occurred overnight at 50°C. The target-probe duplexes were then captured by using magnetic beads and by iteratively washing off unhybridized nucleic acids with increasingly stringent buffers. Enriched libraries were universally re-amplified for 20 cycles using Illumina adapter-specific primers. Normalization and pooling of the enriched libraries were based on quantification using the Quant-iT dsDNA kit (Life Technologies, ON, Canada). Molar quantification of the pooled library was performed using GeneRead Library Quant Kit for Illumina (Qiagen Canada, ON). The pooled library was sequenced as 2×150 nt reads on an Illumina NextSeq 500 instrument, as per the manufacturer’s instructions.

### ONETest: NGS data analysis

Reads from the ONETest libraries were analyzed using an in-house bioinformatics pipeline. The pipeline preprocesses raw NGS reads using a custom C/C++ program (removing adapter sequences, trimming off poor-quality bases of < Q30, and filtering out reads of < 50 nt and reads with low complexity of normalized trimer entropy of < 60, poor mean base quality of < Q27, or percent G of > 40%). Reads were discarded that mapped to the human genome sequence (GRCh38.p13, release 35) using *bowtie2* v2.4.2 ^18^. Then, it aligned the remaining reads to the SARS-CoV-2 Wuhan-Hu-1 reference sequence (MN996528.1) using *bowtie2* (with the settings ‘--very-sensitive-local --score-min G,100,9’), marking duplicate reads using *samtools* v1.11 ^16^. Finally, the pipeline performed iterative comparative assembly (up to five attempts) to reconstruct consensus SARS-CoV-2 genome sequences using *bcftools* v1.11. Nucleotides were called at positions that were covered by ≥ 10 reads (excluding duplicate reads); otherwise, they were masked as Ns. Discounting poor-quality bases of < Q15 and excluding duplicate reads, nucleotide variants were filtered out unless (1) their quality score was ≥ Q15, (2) they were supported by > 1 forward aligned read and > 1 reverse aligned read, (3) they were supported by > 25% of the reads, and (4) the number of variant-supporting reads is ≥ the number of reference-supporting reads; a maximum depth of 30,000 was allowed during pileup. Indels were normalized after calling. The pipeline was implemented in C/C++ and Python using a combination of in-house software and third-party tools, including *Biopython* v1.78 ^19^, *bedtools* v2.29.2 ^20^, *pybedtools* v0.8.1 ^21^, *samtools/bcftools/htslib* v1.11 ^16^, and *Snakemake* v5.26.1 ^22^.

### ONETest: sub-sampling analysis

We sequenced the ONETest libraries at 2.66 million 2×150 nt reads on average, nearly four times as deep as that of the ARTIC libraries (0.63 million 2×150 nt reads on average). To assess whether the observed differences in genome coverage between the ONETest and ARTIC libraries might have resulted from deeper sequencing of the ONETest libraries, we conducted a sub-sampling analysis in which we compared down-sampled ONETest libraries with the full ARTIC libraries. Using *seqtk* v1.3 (https://github/com/lh3/seqtk), we randomly down-sampled (without replacement) the 2×150 nt reads of each ONETest library so that the resulting library had the same number of reads as the matched ARTIC library; each ONETest library was sub-sampled three times in this manner to generate three simulated replicates of the library. Then, we analyzed those sub-sampled reads to determine which bases were poorly covered across the SARS-CoV-2 genome in the simulated ONETest libraries.

### Depth of coverage analysis

Using *bedtools*, we generated depth of sequence coverage profiles for the full ONETest libraries and the sub-sampled ONETest libraries based on *bowtie2* read alignments and the ARTIC libraries based on the *bwa mem* read alignments. For the ONETest libraries, we excluded duplicate reads, but for the ARTIC libraries, we included duplicate reads. Visualization was done in R using *ggplot2* ^23^.

### Lineage analysis

We identified the lineages of SARS-CoV-2 in the samples based on the ONETest and ARTIC consensus sequences using *pangolin* v2.1.10 (https://github.com/cov-lineages/pangolin). This tool assigns SARS-CoV-2 lineages according to a dynamic nomenclature system ^24^.

## DATA AVAILABILITY

The complete or near-complete consensus SARS-CoV-2 genome sequences from the ONETest libraries are available via GISAID (accessions: To be deposited during submission). All de-identified FastQ files (with human reads removed) of the ONETest and ARTIC libraries are publicly available via the NCBI Short Read Archive (BioProject: To be deposited during submission).

## ACKNOWLEDGMENTS

We thank Dr. Mohammad A. Qadir (Fusion Genomics Corp.) for providing guidance throughout this study and constructive feedback on this manuscript and Greg Stazyk (Fusion Genomics Corp.) for setting up the computing infrastructure that enabled this study. We are grateful to Compute Canada and Simon Fraser University for providing the computing resources that facilitated this study. Also, we gratefully acknowledge the support of the staff from the University of Florida Health Shands Hospital Laboratory.

## AUTHOR CONTRIBUTIONS

S. H. Z.: Conceptualization, Methodology, Software, Formal analysis, Investigation, Data curation, Visualization, Writing – original draft preparation, Writing – review and editing.

S. M. A.: Conceptualization, Methodology, Formal analysis, Investigation, Data curation, Project administration, Writing – review and editing.

B. S. K.: Conceptualization, Methodology, Formal analysis, Investigation, Data curation, Project administration, Writing – review and editing.

M. H. L.: Methodology, Software, Formal analysis, Investigation, Writing – review and editing.

J. K.: Methodology, Formal analysis, Data curation, Investigation.

H. D.: Formal analysis, Data curation, Investigation, Writing – review and editing.

H. J. H.: Methodology, Data curation, Resources.

K. H. R.: Conceptualization, Methodology, Investigation, Data curation, Resources, Writing – review and editing.

## SUPPLEMENTARY MATERIAL

**Table S1**. Information about the ONETest and ARTIC libraries and results of SARS-CoV-2 genome sequence analysis. For each sample, the sequence name in GISAID accession, collection date, and sample type, and the SARS-CoV-2 genes (N, RdRp, and E) detected by the OSANG PCR assay are indicated. For each library, the total number of paired-end reads, the number of reads mapped to the SARS-CoV-2 genome (for the ONETest libraries, read pairs were counted, but for the ARTIC libraries, reads were counted), the length of its consensus genome sequence (excluding the Ns at the ends), and mean depth of coverage over the genome (excluding duplicate reads in the ONETest libraries, and including duplicate reads in the ARTIC libraries), the lineage assigned to its consensus sequence using *pangolin* are provided. Abbreviations: NP = nasopharyngeal; ETA = endotracheal aspirates; N/A = not assigned or not available.

**Figure S1.**
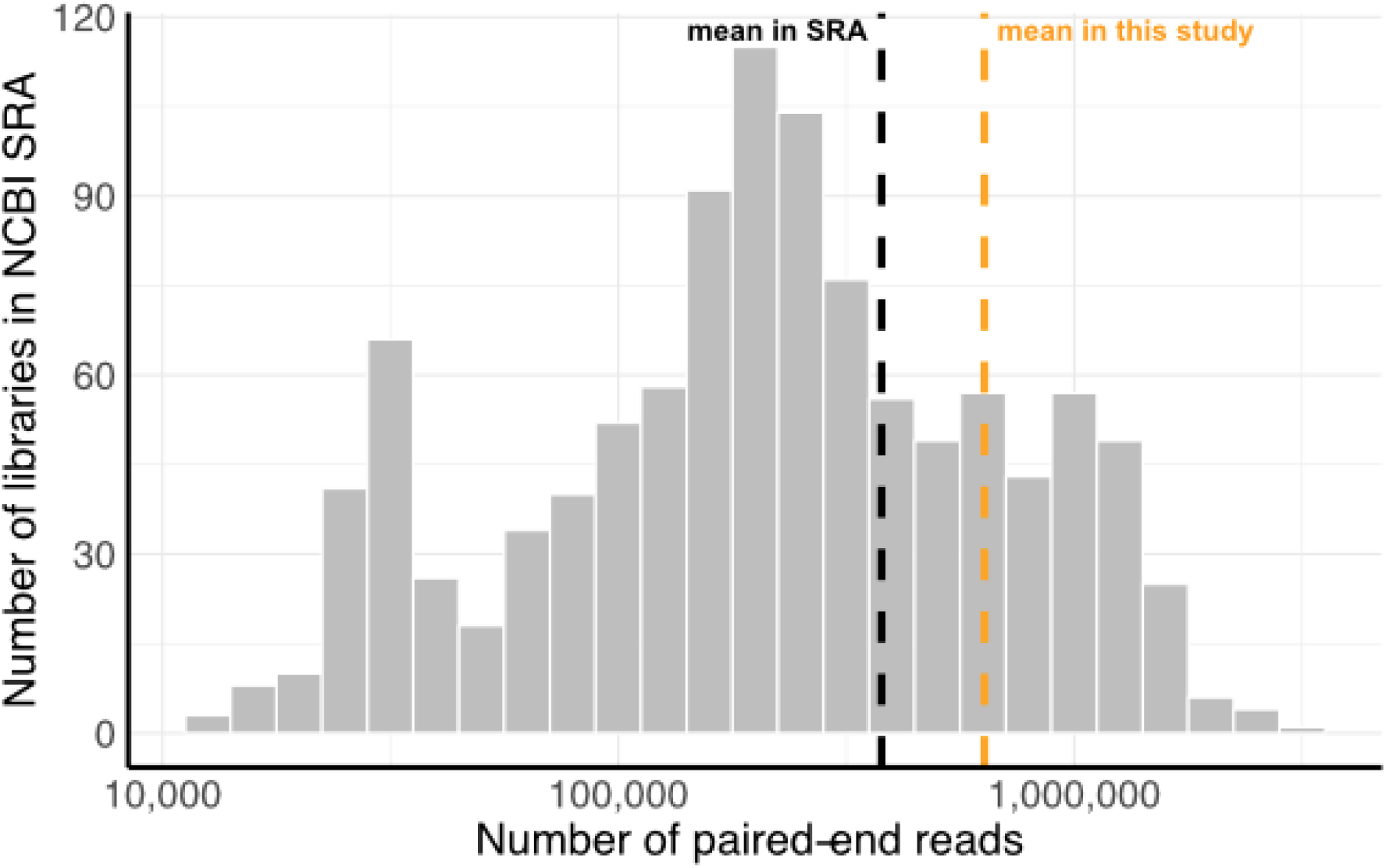
Amount of sequencing in the ARTIC Illumina libraries in the NCBI Short Read Archive. We searched the SRA for 2×150 nt ARTIC Illumina libraries using the query “(((((*Severe acute respiratory syndrome coronavirus 2[Organism]) AND Illumina[Platform]*) *AND PAIRED[Layout]*) *AND 150[ReadLength]*) *AND AMPLICON[Strategy]*) *AND ARTIC*” and then again using the same query except *“149[ReadLength]*” (accessed on Mar. 6, 2021). Ten libraries with < 10,000 paired-end reads were excluded. Also, we excluded entries from SRP287442, which involved sequencing SARS-CoV-2 genomes in cell cultures and mouse models. The vertical black dashed line indicates the mean number of paired-end reads in 1,089 ARTIC libraries in the SRA (0.38 million ± 0.42 million), and the orange line indicates the mean number of paired-end reads in the ARTIC libraries in this study (0.63 million ± 0.30 million).

**Figure S2.**
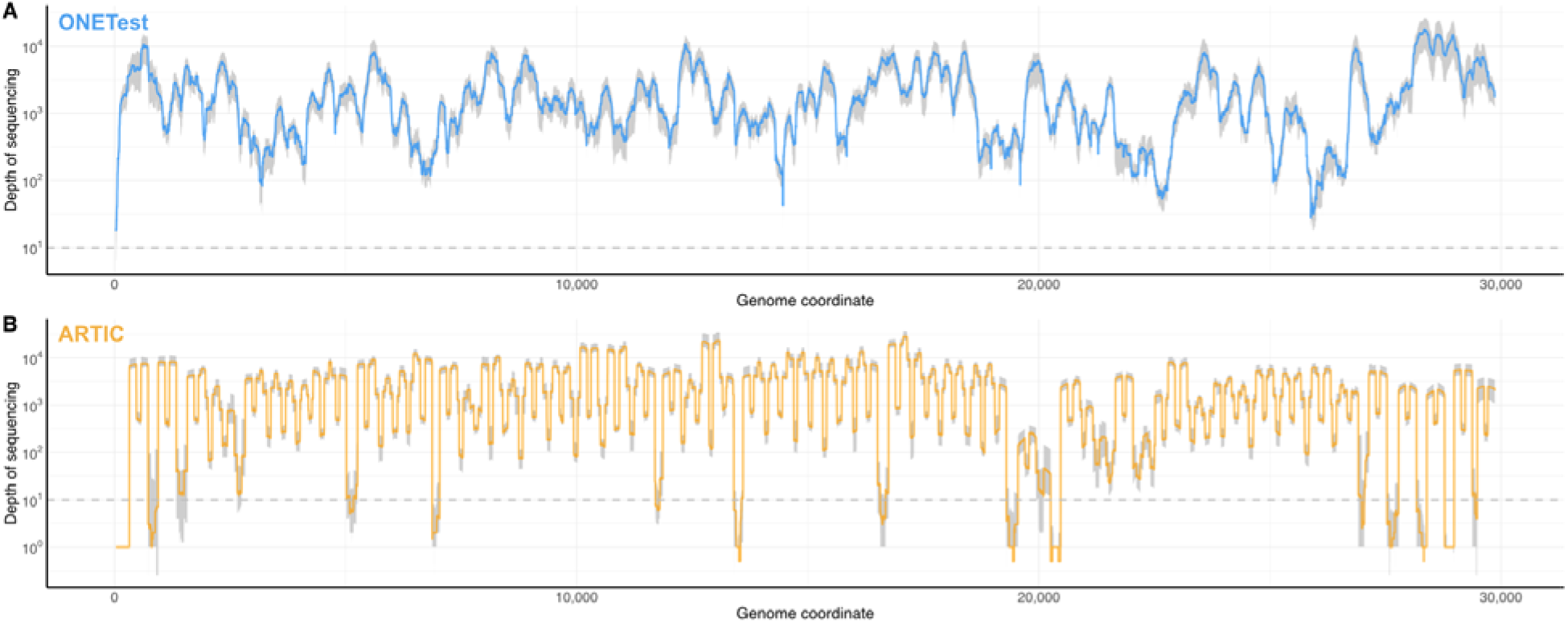
Depth of sequencing coverage over the SARS-CoV-2 genome in the 34 matched pairs of ONETest library (A) and ARTIC library (B) for which lineage could be assigned. Duplicate reads in the ONETest libraries were excluded, and duplicate reads in the ARTIC libraries were included. The y-axis is shown in log10 scale; zeroes were set to one for visualization in log10 scales. The colored line represents the median, and the grey area indicates the 25%-75%tile range. The dashed horizontal line indicates the minimum threshold (≥ 10 depth) to call a base in the consensus genome sequences.

**Figure S3.**
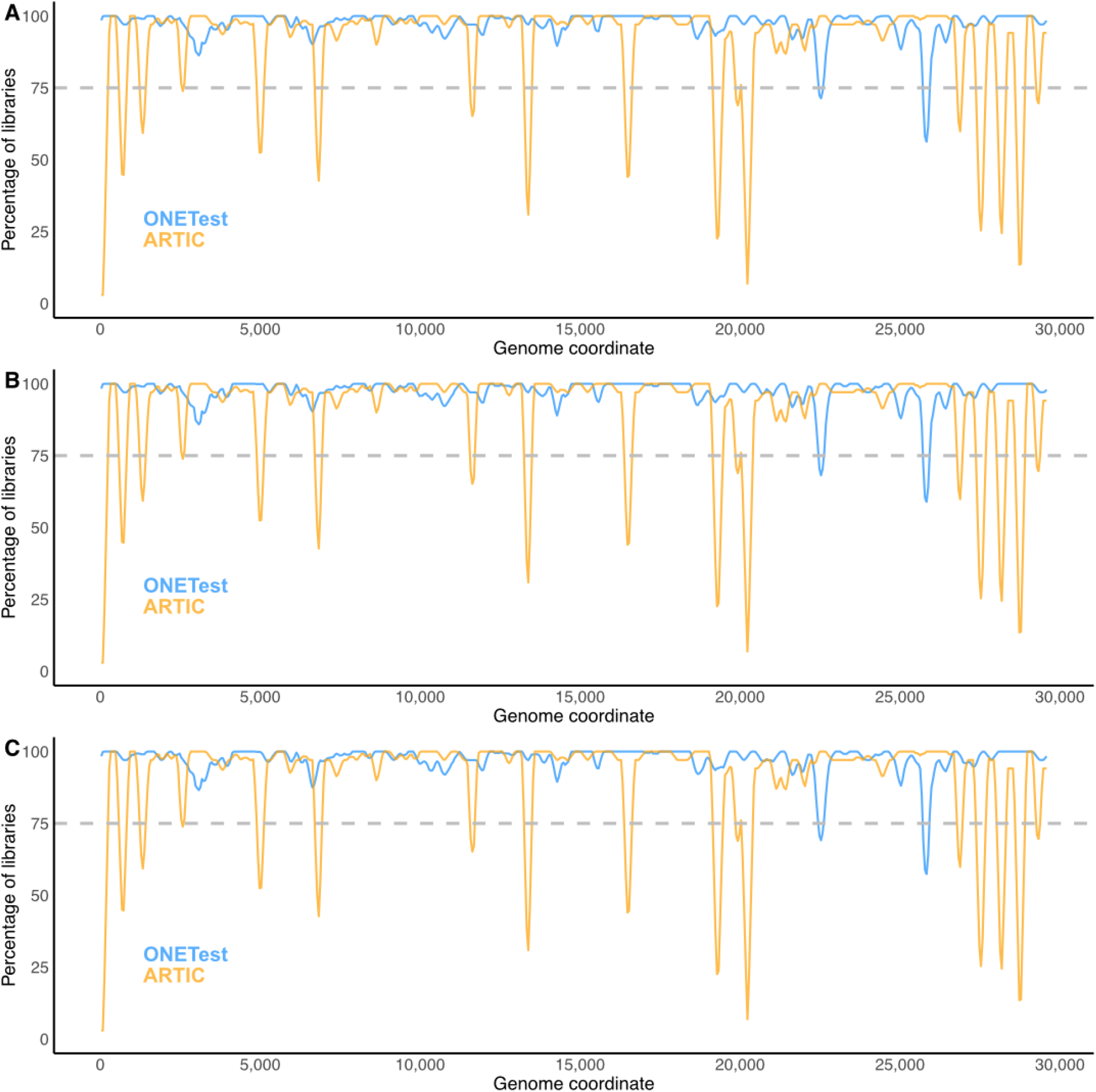
Aggregate summary of sequence coverage over the SARS-CoV-2 genome in the sub-sampled ONETest libraries and the full ARTIC libraries. For each of the 34 samples for which lineage could be assigned to both its ONETest and ARTIC sequences, we randomly sub-sampled its ONETest library three times so that the sub-sampled read sets had the exact number of raw reads as the matched ARTIC library. These data were analyzed the same way as described in **Figure 1**.

**Figure S4.**
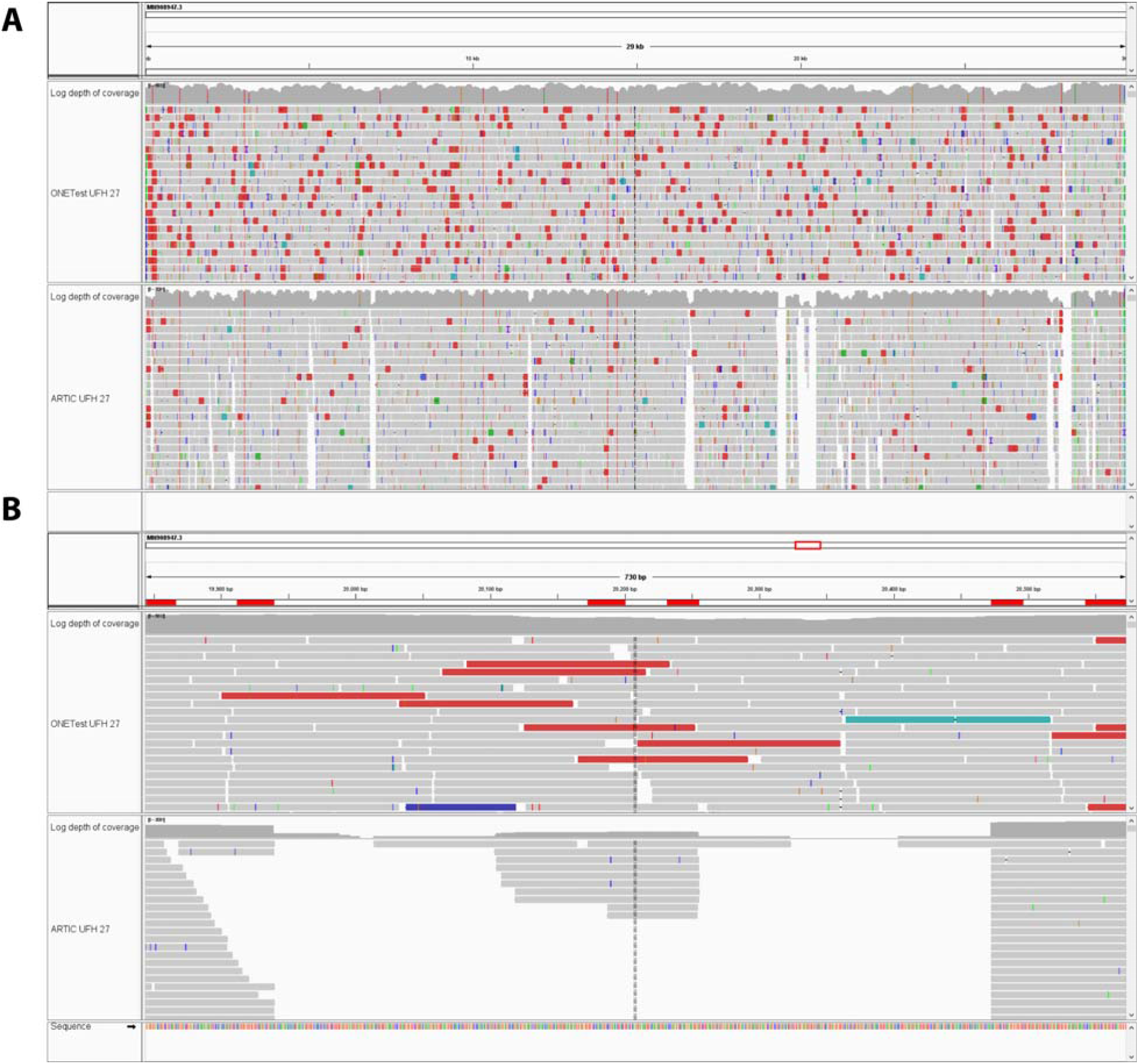
Read alignments over the entire SARS-CoV-2 genome (MN996528.1) (A) and over the 19,844-20,572 region (B) in the ONETest library and the ARTIC library of sample 27. The 19,844-20,572 region is targeted by two ARTIC V3 primer pairs (66_LEFT/66_RIGHT, MN908947.3: 19,844-20,255; 67_LEFT/67_RIGHT, MN908947.3: 20,172-20,572). Visualization was performed using IGV.

